# Identification of tissue-specific and common methylation quantitative trait loci in healthy individuals using MAGAR

**DOI:** 10.1101/2021.05.30.445237

**Authors:** Michael Scherer, Gilles Gasparoni, Souad Rahmouni, Tatiana Shashkova, Marion Arnoux, Edouard Louis, Arina Nostaeva, Diana Avalos, Emmanouil T. Dermitzakis, Yurii S. Aulchenko, Thomas Lengauer, Paul A. Lyons, Michel Georges, Jörn Walter

## Abstract

**Background:** Understanding the influence of genetic variants on DNA methylation is fundamental for the interpretation of epigenomic data in the context of disease. There is a need for systematic approaches not only for determining methylation quantitative trait loci (methQTL) but also for discriminating general from cell-type-specific effects.

**Results:** Here, we present a two-step computational framework *MAGAR*, which fully supports identification of methQTLs from matched genotyping and DNA methylation data, and additionally the identification of quantitative cell-type-specific methQTL effects. In a pilot analysis, we apply *MAGAR* on data in four tissues (ileum, rectum, T-cells, B-cells) from healthy individuals and demonstrate the discrimination of common from cell-type-specific methQTLs. We experimentally validate both types of methQTLs in an independent dataset comprising additional cell types and tissues. Finally, we validate selected methQTLs (*PON1*, *ZNF155*, *NRG2*) by ultra-deep local sequencing. In line with previous reports, we find cell-type-specific methQTLs to be preferentially located in enhancer elements.

**Conclusions:** Our analysis demonstrates that a systematic analysis of methQTLs provides important new insights on the influences of genetic variants to cell-type-specific epigenomic variation.

## Background

Epigenetic mechanisms, including histone modifications, small RNAs, and DNA methylation, regulate gene expression in a tissue- and cell-type-specific manner [1]. DNA methylation is a critical player in such epigenetic gene regulation that has been implicated in various biological processes including X-chromosomal inactivation [2], genomic imprinting [3], and allele-specific expression [4,5]. DNA methylation has been shown to be highly cell-type-specific and can be used to reliably estimate the proportions of different cell types in mixed cell samples such as blood or tissues [6,7]. The DNA methylation state of a defined subset of CpGs in the human genome can be measured reliably across many samples using the Illumina Infinium microarray technologies allowing to perform epigenomic association studies (EWAS).

DNA methylation can be affected by aging [8], sex, and a range of environmental exposures [9,10]. Additionally, donor genotype has a strong influence on the global DNA methylation state (methylome), especially when a genetic alteration, such as a single nucleotide polymorphism (SNP), occurs at a CpG site. Since bisulfite-based methods can generate unclear and uninterpretable data at annotated or predicted SNPs located at CpG dinucleotides, such positions are typically removed from the analysis of DNA methylation data [11].

However, additional genetic effects that are not located in the CpG site but in genetic variants distant to the analyzed CpG can influence its DNA methylation state. Such variants influencing DNA methylation states are referred to as methylation quantitative trait loci (methQTL). These associations can range from distances of a few bases to several megabases resulting in long-range interactions [12,13]. The definition of proximal methQTLs varies from 500 kb to 2 mb distance between the CpG and the SNP [12–14]. MethQTLs co-localize with genetic variants associated with diseases and donor phenotypes (GWAS hits) including obstructive pulmonary disease [14], prostate cancer risk [15], osteoarthritis [16], immune-mediated disease [17], asthma [18], and smoking [19]. Furthermore, combining methQTLs with expression QTLs (eQTLs) enables the investigation of associations between DNA methylation and gene expression changes [20–22].

However, so far not much emphasis has been put into analyses to investigate if and how often methQTLs affect DNA methylation in a tissue- or cell-type-specific manner. An earlier study used cultured cells including fibroblasts, T-cells, and lymphoblastoid cell lines to determine largely tissue-independent methQTLs. The authors reported that the association of methQTLs with changes in gene expression was rather cell-type-specific [23] in line with recently identified cell-type-specific eQTLs [24]. Other studies analyzing primary human cells rather reported largely cell-type-independent eQTLs [25]. One problem which may have contributed to the current mixed view on the distribution of methQTLs is that methQTLs are typically determined using statistical models and tools that have been developed for eQTL analysis (e.g., *Matrix-eQTL* [26], *fastQTL* [27], or *GEM* [28]). Without the consideration of the specific properties of DNA methylation data including the correlation of DNA methylation states of neighboring CpGs such approaches may lead to substantial biases in the calling and interpretation of methQTLs.

To alleviate this problem, we present “Methylation-Aware Genotype Association in R” (*MAGAR*) - a novel computational pipeline that performs methQTL analysis. *MAGAR* defines clusters of neighboring CpGs according to their shared behavior across samples to represent DNA methylation haplotypes and performs methQTL analysis for each of the correlation blocks independently. *MAGAR* has been implemented as an R-package and utilizes existing tools such as *fastQTL* [27], *RnBeads* [29,30], and *PLINK* [31]. Using *MAGAR*, we investigated sorted blood cell types (T-cells, B-cells) and composite bowel tissues (ileum, rectum) of healthy individuals. The identified methQTLs were analyzed for cell-type-specific effects using colocalization analysis, which showed that we could discern tissue-specific from common methQTLs. Finally, we validated and reproduced our findings in additional samples and in data from two published methQTL studies.

## Results

### Strong cell-type-specific DNA methylation signals identified in bowel biopsies and purified blood cell types

The data set that we used for the discovery of methQTLs comprised 409 samples from ileum (IL, n=98) and rectum (RE, n=95) tissue biopsies and the two FACS sorted blood cell types CD4-positive T-cells (n=119) and CD19-positive B-cells (n=97). For 29 individuals DNA methylation data was available for all four tissues/cell types within this discovery data set (**Supplementary Figure 1**). Average DNA methylation levels across all CpGs in genome-wide five-kb bins revealed a strong cell-type-specific signal that discriminates the blood cell types from the biopsies. Overall, the tissue biopsies exhibited an enhanced variation in comparison to the purified blood cell types indicating that increased cell-type heterogeneity goes along with a higher variation of DNA methylation patterns both on genome-wide bins and on the single-CpG level (**Figure 1A, B**). To better understand the origins of cellular heterogeneity within the biopsy samples we estimated the overall immune-cell content of a sample using the LUMP algorithm [32] (**Figure 1C**). While LUMP estimates were uniformly close to one for the two blood cell types as expected, they substantially varied across the biopsy samples. In line with previous reports [33], significantly higher immune cell content was observed in ileal compared to rectal samples.

**Figure 1:**
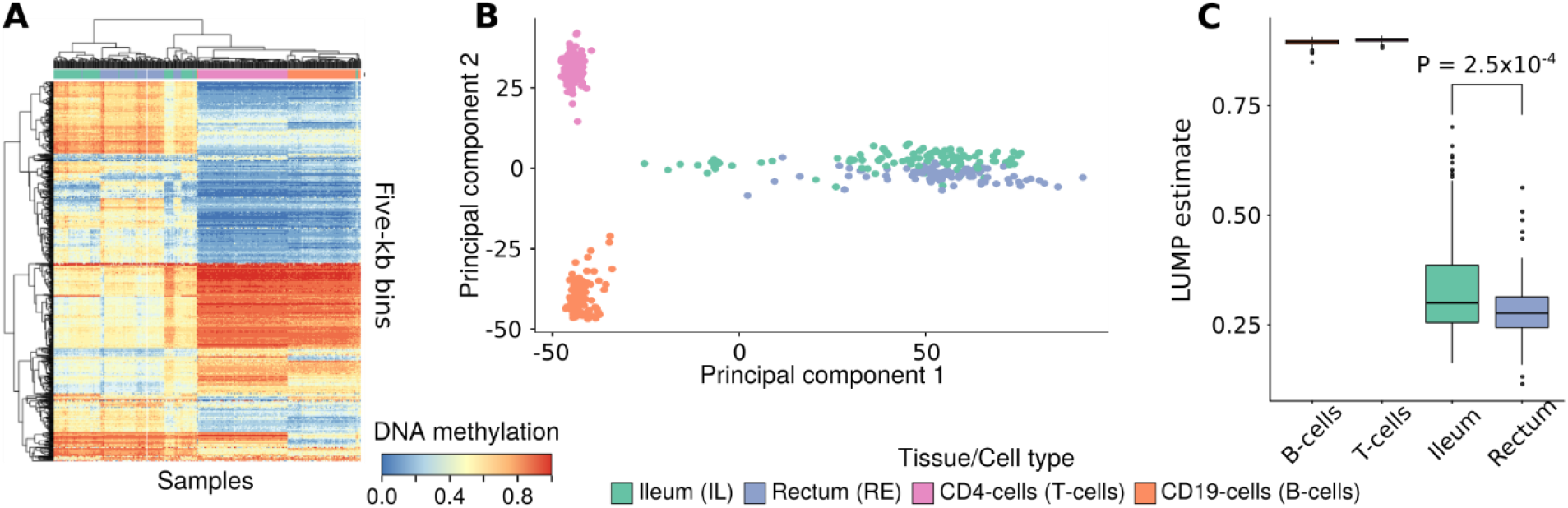
Cell-type-specific DNA methylation patterns in the discovery data set. **A**: Heatmap (blue low, red high DNA methylation levels) of the 1,000 most variably methylated genome-wide bins of size 5 kb. Hierarchical clustering of samples and bins was performed using Euclidean distance and complete linkage. **B:** PCA plot of genome-wide DNA methylation data at the single-CpG level. The first two principal components are displayed. **C:** Boxplots depicting the distributions of LUMP estimates for the overall immune cell content of the different cell types/tissues. The P-value was computed using a two-sided t-test.

### *MAGAR* facilitates the analysis of genome-wide methQTL effects

Understanding the relationship between DNA methylation and genetic variants can help to illuminate the association of genetic alterations with diseases and changes in gene expression. Thus, we are interested in defining statistically significant associations between DNA methylation and genotyping data. We call genetic variants that are associated with DNA methylation methQTLs. To alleviate the methQTL identification process, we developed the new R-based framework *MAGAR* (**M**ethylation-**A**ware **G**enotype **A**ssociation in **R**) that provides a comprehensive suite of tools enabling methQTL analysis leveraging the correlation of DNA methylation states of neighboring CpGs (**Figure 2A**). Notably, *MAGAR* is the first package that performs data processing of raw (i.e, IDAT files) DNA methylation and genotyping data before returning data formatted for methQTL analysis.

**Figure 2:**
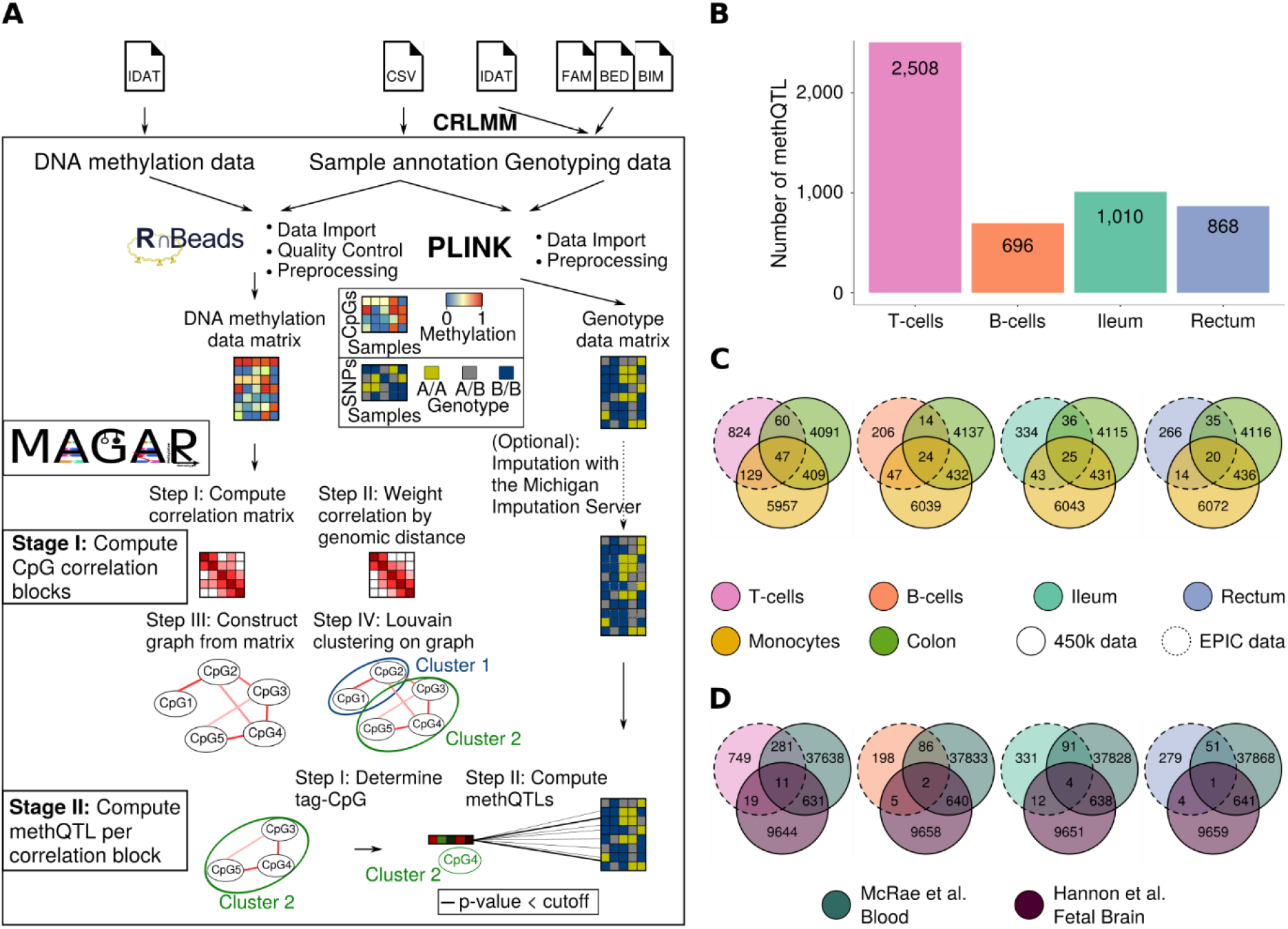
Overview of MAGAR and methQTL results. ***A**: MAGAR* is an R-package utilizing a two-stage protocol. After data import via established software packages, CpGs are clustered into CpG-correlation blocks in a four-step procedure. In the second stage, methQTLs are called for each correlation block separately. **B:** Number of methQTLs identified by *MAGAR* for T-cells, B-cells, ileum, and rectum samples. Overlap between the methQTLs identified per tissue/cell type with methQTLs identified in the validation cohort (**C**) and in published methQTLs from blood [12] and fetal brain samples [37] (**D**). The methQTLs were reduced to those methQTLs affecting CpGs present on the 450k microarray.

In the first phase of *MAGAR*, raw data is converted and processed using the established software packages *RnBeads* [29,30], *PLINK* [31] and *CRLMM* [34,35]. The processing includes data filtering of CpGs and SNPs according to quality criteria (see Methods for details). The second phase of the package – the methQTL calling – has been implemented as a two-stage workflow as follows: Initially, CpGs that exhibit high correlations of methylation states across the samples are clustered into CpG correlation blocks. *MAGAR* takes into consideration that the DNA methylation states of neighboring CpGs in the same functional or regulatory unit are usually highly correlated [36]. This assumption implies that one may not need to inspect each CpG. In fact, doing so would generate many redundant methQTLs. In *MAGAR* we therefore group neighboring, highly correlated CpGs into correlation blocks. In the second stage of the process, methQTLs are determined individually for each of the CpG correlation blocks. To this end, for each correlation block, *MAGAR* determines a tag-CpG representing this block and determines statistically significant associations for each of the tag-CpGs with all SNPs within a specified genomic distance (in this instance 500 kb up- and downstream). This methQTL calling can either be performed using univariate, linear least squares or by the approach implemented in *fastQTL* [27]. The f*astQTL* software computes correlations between DNA methylation states and SNP genotypes and uses a permutation scheme to address the multiple testing problem. *MAGAR* provides various options for modifying the analysis, including options for defining the CpG clustering, for defining the tag-CpG per correlation block, and for the methQTL calling approaches to be employed (linear modeling or *fastQTL*). Reasonable default values for the parameters were selected using simulation experiments (**Supplementary Text, Supplementary Figure 2**). *MAGAR* returns a list of associations and corresponding statistics, which can be filtered further by the user to define methQTLs or which can be used in downstream analyses. In the analysis presented here*, MAGAR*’s output was used as input to colocalization analysis for defining tissue specificity.

Using *MAGAR*, we analyzed the ileal, rectal, T-cell, and B-cell methylation data (659,464 CpGs) jointly with genotype data from 5,436,098 SNPs and calculated methQTL statistics for each cell type/tissue independently. To determine significant methQTLs, we selected a Bonferroni-corrected genome-wide P-value cutoff of 8.65×10^−11^ (see Methods for details). As a result, we found 696, 2,508, 1,010, and 868 methQTLs for CD19+ B-cells, CD4+ T-cells, ileal, and rectal biopsies, respectively (**Figure 2B**, **Supplementary Table 1**). To validate the methQTLs, we used additional samples from monocytes and transverse colon from the same cohort (**Supplementary Figure 3**). Additionally, we obtained published methQTLs from two studies (blood [12] and fetal brain [37]) and compared them with the identified methQTLs. Note that the validation cohort and the published studies used DNA methylation data generated using the 450k microarray, which comprises fewer CpG sites than the EPIC array. Thus, we excluded those methQTLs from the comparison that associated with a CpG site that is exclusively present on the EPIC array. We identified some of the methQTLs found in the discovery cohort using a different, validation P-value cutoff (see Methods) in the validation cohort (**Figure 2C**) and in the published data (**Figure 2D**). Notably, the overlap between the identified and the published methQTLs was significantly higher than expected by chance (**Supplementary Table 2**). As expected, the overlap of the methQTLs identified in B- and T-cells with the methQTLs identified using whole blood was higher than with those identified in fetal brain samples (**Figure 2D**).

### Colocalization analysis identifies common methQTLs

Next we applied colocalization analysis that uses summary statistics from two association studies (here methQTLs in two different tissues) to determine if an association of two traits (here CpG methylation states) to the same genetic region is significant and is likely to be caused by the same pleiotropic genetic variant. Colocalization was examined using Summary-data-based Mendelian Randomization (SMR) analysis followed by the Heterogeneity in Dependent Instruments (HEIDI) test [38]. The SMR test indicates whether the two traits are significantly associated with the same locus and the HEIDI test interrogates whether the data are compatible with the hypothesis that both traits are affected by the same underlying functional SNP.

We only included methQTLs in the analysis that were significant at P-value lower than 8.65×10^−11^ in at least one tissue. The analysis is anchored at the tissue where the methQTL exhibited a significant association and the methQTL statistics were compared with those in the other tissues. In total, 4,253 colocalization tests were performed (**Supplementary Table 3**) based on the number of significant methQTLs. We defined those methQTLs as shared between two tissues/cell types that had an FDR-adjusted P-value of the SMR test lower than 0.05 and that had a HEIDI test nominal P-value larger than 0.05 (**Supplementary Table 3**). These methQTLs are likely driven by the same genetic variant and the shared association is likely caused by a single pleiotropic variant rather than two linked variants. Colocalization analysis was conducted for all pairs of cell types/tissues (**Figure 3A**) and we define three classes of methQTLs:

1. Common methQTLs are shared across all investigated pairs of tissues/cell types according to the colocalization analysis <u>and</u> pass the methQTL P-value cutoff 8.65×10^−11^ in all tissues
2. Shared methQTLs are shared across all the investigated pairs of tissues/cell types according to the colocalization analysis
3. Tissue-specific methQTLs are only present in one of the tissues/cell types and not shared in any pairwise comparison according to the colocalization analysis

**Figure 3:**
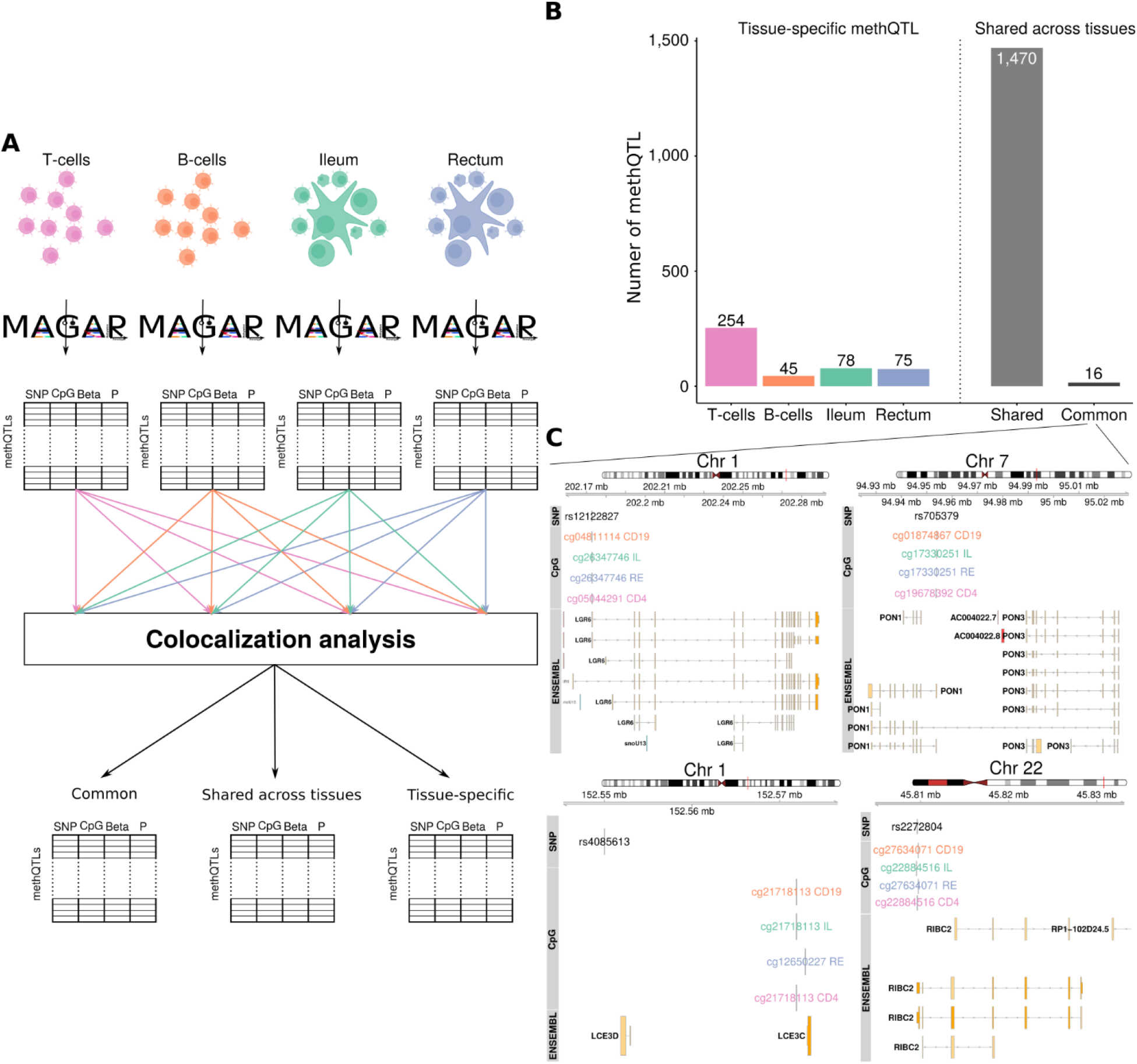
Common and tissue-specific methQTLs identified through colocalization analysis. **A**: To define tissue specificity, we employed MAGAR on the four tissues/cell types independently to obtain methQTL statistics. These were used in pairwise colocalization analyses to define common and tissue-specific methQTL, as well as methQTLs shared across several tissues. **B:** Number of tissue-specific methQTLs per tissue and methQTLs shared across different tissues according to the colocalization analysis. Common methQTLs were shared according to the colocalization analysis and had methQTL P-values below the cutoff in all tissues. **C:** Examples of four common methQTLs located in vicinity to PON1, LGR6, LCE3D, and RIBC2.

We found that 16 methQTLs were shared across all of the pairwise comparisons and lay below the methQTL P-value cutoff of 8.65×10^−11^ in all tissues and are thus common methQTLs (**Figure 3B, Supplementary Table 4**). The common methQTLs included well established methQTLs and eQTLs, such as the ones present in the *PON1* [39], *LGR6* [40], and *RIBC2* [41] loci (**Figure 3C**). We found substantially more methQTLs shared across different tissues than tissue-specific methQTLs. Most (254) tissue-specific methQTLs were exclusively found in CD4+ T-cells (**Figure 3B, Supplementary Table 5**), and similar numbers of tissue-specific methQTLs (78, 75) were identified for ileal and rectal biopsies, respectively. Due to the definition above, common methQTLs are a subset of the shared methQTLs.

We used the validation cohort to validate the identified common and shared methQTLs further. Notably, the validation cohort samples were assayed using the 450k array and only 10 (of 16) and 689 (of 1,470), respectively, of the common and shared methQTLs associated with a CpG present on the 450k array. We found that most of the common (9/10, Fisher test P-value: 1.6×10^−4^) and some of the shared QTLs (178/689, Fisher test P-value: 1) were also present in at least one of the two tissues (**Supplementary Figure 4A,B**). Additionally, four of the 10 overlapping common methQTLs (rs2272804, rs705379, rs55901738, rs10021193) were also identified in an independent study of blood samples [12] (**Supplementary Figure 4C**).

### Common methQTL at *PON1* locus identified in independent samples using ultra-deep bisulfite sequencing

To rule out potential technology-dependent artifacts, we used local deep amplicon sequencing for the validation of a common methQTL. We selected the methQTL at the *PON1* locus (comprising rs705379, cg19678392, cg17330251, and cg01874867), since both the SNP and the CpGs could be included into a single amplicon of size 462 base pairs. Thus, we were able to capture the genotype of the SNP and the DNA methylation state of multiple CpGs simultaneously. Notably, we associated the genotype with the CpG methylation state at the single-molecule level, since each sequencing read represents a single molecule. The results indicated a strong relationship between the genotype of rs705379 and the CpG methylation state at all CpGs present in the amplicon, while the effect was stronger in those CpGs that were closer to the SNP (**Figure 4A**). In this setting, the A genotype was associated with a high DNA methylation state of more than 50%, while the G genotype leads to a decrease of the methylation level below 25% for some CpGs (**Figure 4B**). Notably, there was no one-to-one relationship between the genotype and the DNA methylation state and G genotypes co-occurred with methylated CpGs and A genotypes with unmethylated CpGs at the single-molecule level. The effect of the SNP on the DNA methylation state was consistent across all samples within a genotype, and the standard deviations across the samples within the different genotype groups were comparable (T-cells: AA: 0.054, AG: 0.046, GG: 0.058; B-cells: AA: 0.042, AG: 0.058, GG: 0.06, **Figure 4C**). Notably, rs705379 had a high minor-allele frequency of 0.46 for the B-cell and 0.47 for the T-cell samples in our cohort. To further investigate whether the effects that we detected are also present for methQTLs beyond the 16 common methQTLs, we constructed two additional amplicons to capture the methQTLs shared across different cell types/tissues at the *ZNF155* (**Supplementary Figure 5**) and *NRG2* (**Supplementary Figure 6**) loci. In accordance with the results obtained in the *PON1* amplicon, we found a strong association of the genotype with DNA methylation states.

**Figure 4:**
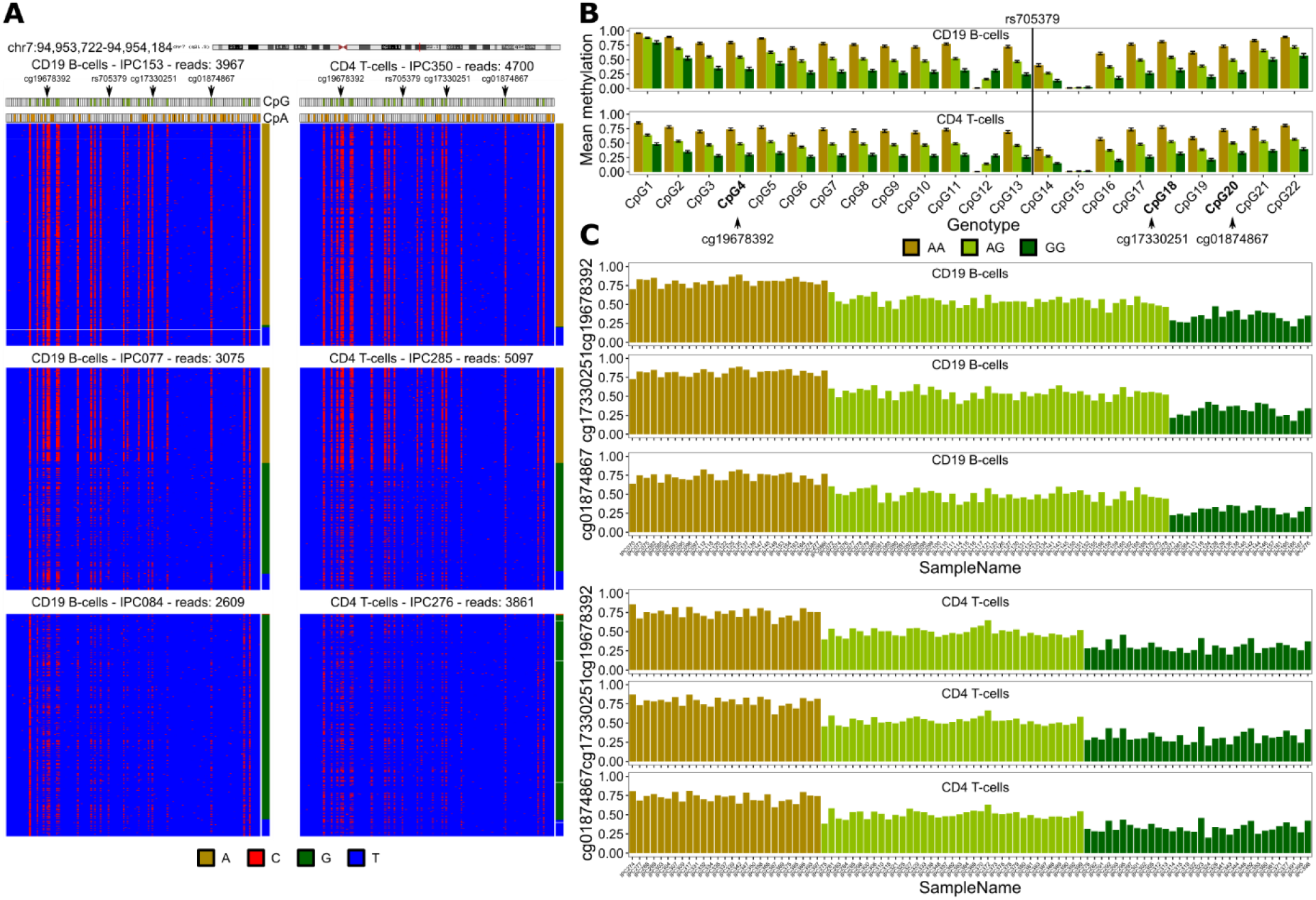
Validation of methQTL at PON1 locus using ultra-deep bisulfite sequencing. **A:** Bisulfite sequencing read pattern maps for three individuals with genotypes homozygous for the reference allele (AA), heterozygous (AG), or homozygous for the alternative allele (GG) for B-cells and T-cells, respectively. Each line is a sequencing read, where the red color indicates a cytosine, i.e., a methylated cytosine before bisulfite conversion, and blue a thymine, i.e., an unmethylated cytosine before bisulfite conversion. All cytosines within the amplicon are shown in the pattern map and the CpG and CpA dinucleotides are marked. The genotype at rs705379 per sequencing read is indicated on the right. Shown is the common methQTL at the PON1 locus at chr7:94,953,722-94,954,184 (hg19). **B:** Average DNA methylation levels across all samples of the same genotype and standard deviations across the samples. The barplots are shown for all 22 CpGs present in the amplicon. **C:** Average DNA methylation levels across all sequencing reads per sample for the three CpGs that were associated with the SNP genotype in the microarray data analysis for B-cells and T-cells.

### Tissue-specific methQTLs are preferentially located in proximal enhancer elements

To determine characteristic properties of tissue-specific methQTLs, we compared all 452 tissue-specific methQTLs with 1,470 methQTLs shared across multiple tissues (**Supplementary Table 6**). While the distance between the CpG and the SNP that significantly correlates with the DNA methylation state was not different in the two classes of methQTLs, we found both stronger effects on the DNA methylation state with respect to effect size and lower P-values for the shared methQTLs than for the tissue-specific methQTLs (**Figure 5A**). To determine whether the CpGs or the SNPs of the shared and cell-type-specific methQTLs are preferentially located in particular functional regions of the genome, we performed enrichment analysis for various functional annotations such as gene promoters and proximal enhancers. We found that methQTL SNPs were depleted in regions of active transcription such as transcriptional start sites (TSS) and gene bodies for the shared methQTLs (**Figure 5B**). No significant enrichment of a functional category was detected for the shared methQTLs. In contrast, the tissue-specific methQTLs were preferentially located in proximal enhancer elements according to the Ensembl Regulatory Build [42] further pointing toward the important regulatory role of enhancers in establishing cellular identity. Further indication for this hypothesis was obtained by the LOLA [43] enrichment of tissue-specific methQTLs in enhancer elements and transcription factor binding sites indicating an enhancer element in B-cells and in the B-lymphocyte cell line GM12878 (**Figure 5C**). Analogously, we associated the tissue-specific and shared methQTL SNPs and CpGs with overlapping gene bodies. For those overlapping genes, we performed Gene Ontology (GO) enrichment analysis [44] and detected an enrichment of the shared methQTLs towards the biological process “cell development” (P-value=0.0069, **Supplementary Table 7**).

**Figure 5:**
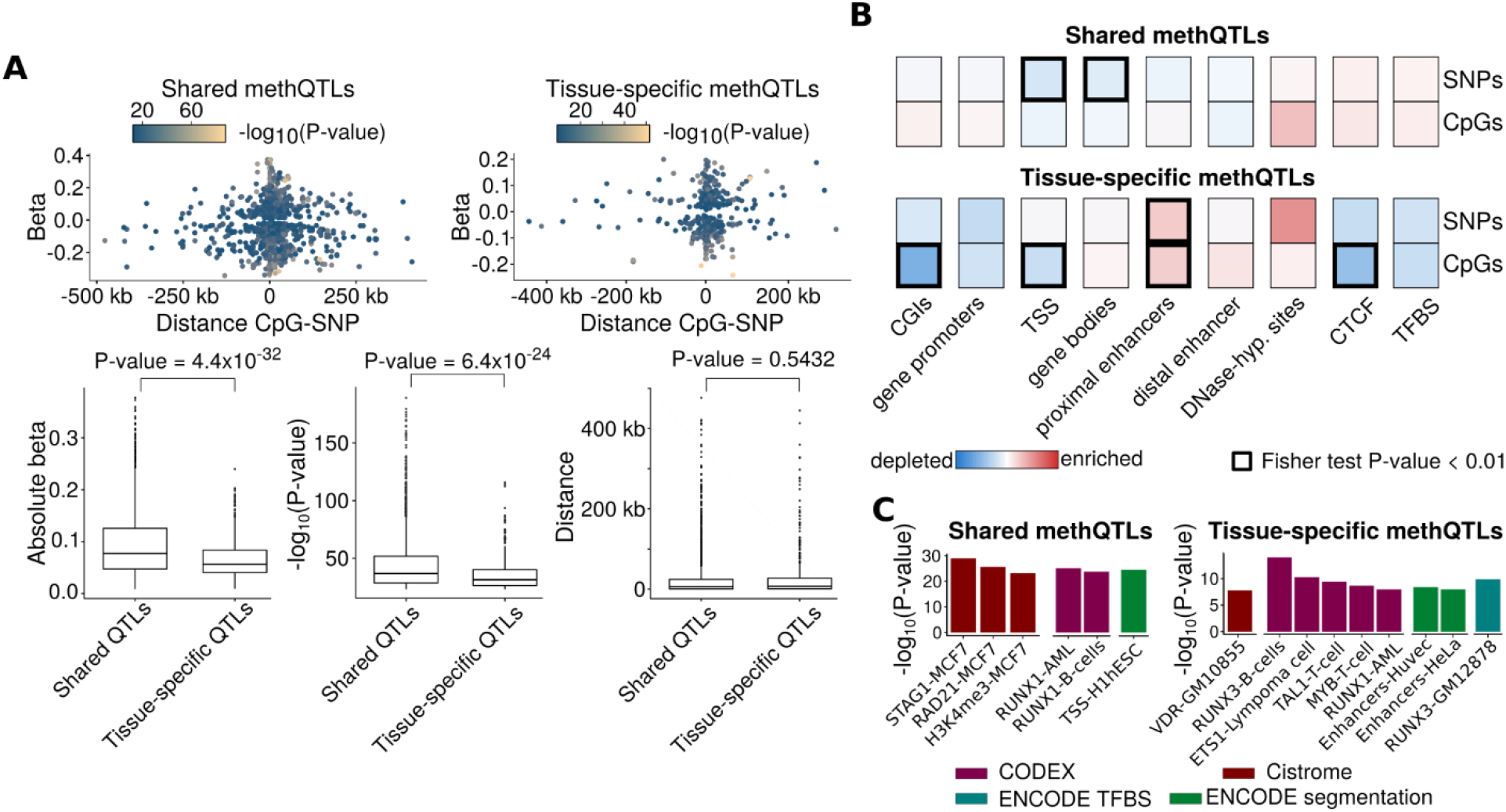
Properties of methQTLs shared across the tissues and tissue-specific methQTLs. **A:** Distance between the CpG and the SNP, the effect size (slope of the regression) of the methQTL, and the negative common logarithm of the methQTL P-value are visualized. MethQTLs were classified as either shared or tissue-specific. **B:** Enrichment analysis of shared (top) or tissue-specific methQTLs (bottom) in different functional annotations of the genome. Visualized is the common logarithm of the odds ratio and the associated Fisher exact-test P-value was computed. P-values below 0.01 are indicated by a bold outline. **C:** LOLA [43] enrichment analysis of the methQTL SNPs for the shared and tissue-specific methQTLs, respectively. ESC=embryonic stem cell, AML=acute myeloid leukemia.

We aimed to validate the tissue-specific methQTLs in the validation cohort and in independent studies. While some of the ileum- and rectum-specific methQTLs identified earlier were present in the transverse colon samples, only two of them were present (at P-value cutoff 9.84×10^−6^) in the monocytes. Similarly, two of the T-cell-specific methQTLs were also found in transverse colon. However, more (seven for T-cells, one for B-cells) were found in the CD14-positive monocytes (**Supplementary Figure 4A**). To validate whether T-cell- and B-cell-specific methQTLs actually capture effects specific to blood cell types, we compared the methQTL effect sizes in the monocytes and in transverse colon. We detected significantly higher effect sizes for the T-cell-specific methQTLs in the monocytes in comparison to transverse colon (**Supplementary Figure 4B**). Notably, not all methQTLs detected in the discovery cohort could be found in the validation cohort, since the latter has been assayed using the Infinium 450k technology. Similarly, more of the T- and B-cell-specific methQTLs were present in the methQTL study on blood samples in comparison to fetal brain samples (**Supplementary Figure 4C**).

## Discussion

Patient-stratification according to mutational signatures, i.e., genotype-based markers, is already well-accepted in the clinic [45]. Recently, DNA methylation-based biomarkers are also becoming relevant in a clinical setting [46] and may contribute to clinical decision making. The relationship between genotype and DNA methylation variation is only just beginning to be understood. As a first step towards the joint characterization of DNA methylation patterns and genotypes, methylation quantitative trait loci (methQTL) have been identified in healthy individuals. To facilitate standardized analyses of DNA methylation and genotyping data, we developed the R-package *MAGAR* that supports processing of raw data and integrates with established bioinformatic tools. *MAGAR* is the first package providing a start-to-finish workflow for microarray-based methQTL studies and supports bisulfite sequencing data, without specifically using the information on co-methylation of neighboring CpGs present in the sequencing reads. For bisulfite sequencing data, specialized methods are available such as *IMAGE* [47]. Notably, *MAGAR* performs methQTL analysis while accounting for the correlation structure of neighboring CpGs and is a first step toward associating genetic haplotypes with DNA methylation haplotypes. Grouping together CpGs into clusters is an approach that has also been used earlier [48,49] in contexts different from methQTL analysis. The earlier approaches to group CpGs into correlation blocks however either do not take into account the genomic distance between two CpGs or are restricted to either microarray or bisulfite sequencing data.

It remains elusive whether methQTLs are inherently cell-type-specific or tissue-independent. In this study, we systematically investigated cell-type specificity of methQTLs in sorted blood cell types (CD19+ B-cells, CD4+ T-cells) and bowel biopsies (ileum, rectum). We found fewer tissue-specific methQTLs than methQTLs that were shared across tissues. We validated tissue-specificity in additional CD14+ monocyte and transverse colon samples. Since DNA methylation is a cell-type-specific epigenetic mark, it is likely that methQTLs are also cell-type specific. It remains to be shown whether these cell-type-specific methQTLs preferentially co-occur with other cell-type-specific epigenetic marks such as open chromatin or histone modifications. Previous methQTL studies [12,37] identified a partially overlapping list of methQTLs, some of which were also detected in this study. Notably, the previous studies used a different strategy for identifying methQTLs (*Merlin* [50] in the blood study and *Matrix-eQTL* [26] in the fetal brain samples). While these strategies do not account for the properties of DNA methylation data, we found a substantial overlap with the methQTLs that we identified.

We found that cell-type-specific methQTLs were preferentially located in enhancer elements, which further emphasizes the importance of enhancers to establish cellular identity. However, methQTL effects were weaker in cell-type-specific methQTLs compared to those shared across different cell types. It remains to be shown how methQTLs affect gene expression states in our samples. In subsequent analyses, the overlap between methQTLs and eQTLs can be explored to further understand the relationship between genome, epigenome, and transcriptome. Since the cell-type-specific methQTLs were associated with the CpG methylation states to a lower extent than shared methQTLs, cell-type-specific methQTLs could modulate transcript abundance in a more fine-grained manner. We would also like to point out that this observation may be due to technical rather than biological reasons. Using colocalization analysis for determining shared effects of methQTLs across tissues, a bias towards stronger effects can be introduced. Since we define tissue-specific methQTLs as those that are not shared according to the colocalization analysis, they could be weaker than the shared ones by definition.

There are some aspects of methQTLs, which remain to be investigated. It would be relevant to study cell-type-specificity of methQTLs in purified cell types outside of the hematopoietic system, such as in neurons, epithelial cells, and hepatocytes. To that end, the identified common methQTLs could be further validated to determine whether they are truly tissue- and cell-type-independent. Furthermore, *MAGAR* groups together CpGs into CpG correlation blocks, which reduces the number of redundant methQTL interactions detected. However, methQTLs affecting single CpGs may be missed using this method. It is well-established that genetic associations with a disease (GWAS hits) are preferentially located in non-coding regions of the genome [51]. The functional impact of such genetic variants, which can be modulated by QTLs (methQTLs, eQTLs), remains to be investigated. Additionally, DNA methylation data can be used to reliably estimate the proportions of different cell types in the samples, either using a reference data set [6] or in a reference-free way [52]; an analysis strategy known as deconvolution. Given the cell-type specificity of a subset of methQTLs identified within this study, a combination of DNA methylation-based deconvolution and identification of methQTLs could be implemented similarly to transcriptome-based approaches [24,53]. By using such a method, it will be possible to investigate methQTL effects in bulk tissues without considering cell-type-specific signals. Preferably, novel analysis methods, such as colocalization analysis and the integration of methQTL and DNA methylation-based deconvolution, are implemented in an easy-to-use software package such as *MAGAR*. To overcome the issue of cell-type specificity, DNA methylation can be assayed at the single-cell level and associated with genotype information from the same cell. Alternatively, more readily accessible single-cell RNA-seq datasets can be jointly analyzed with bulk methQTL studies to understand gene regulation at the single-cell level. Finally, long-read sequencing allows for simultaneously profiling of the genotype and DNA methylation state of the same molecule over distances up to 10 kb, which enables associating genetic haplotypes with DNA methylation haplotypes.

## Conclusions

In summary, the relationships between genetic and epigenetic variations are currently underexplored. To facilitate the joint analysis of genotype and DNA methylation data, we present *MAGAR* as a novel software tool that accounts for the properties of DNA methylation data. In combination with colocalization analysis, we identified tissue-specific and common methQTLs with unique biological properties and genomic location. Tissue-specific and shared methQTLs identified using *MAGAR* were validated in both independent samples and were verified using an alternative local deep sequencing approach.

## Methods

### *MAGAR* R-package

#### *MAGAR* package overview

We developed “**M**ethylation-**A**ware **G**enotype **A**ssociation in **R**” (*MAGAR)* as a new computational framework to determine methQTLs from DNA methylation and genotyping data. *MAGAR* supports both sequencing-based assays including whole-genome (bisulfite) sequencing and microarray-based data. It is the first computational framework for performing methQTL analysis starting from raw DNA methylation and genotyping microarray data. The pipeline implemented within *MAGAR* comprises the following phases:

i. **Data import and preprocessing** using established software packages such as *PLINK* [31], *RnBeads* [29,30], and *CRLMM* [34,35]. Additional modules for quality control and standard processing using these packages are available to the user. *MAGAR* supports automated imputation using the Michigan Imputation Server [54].
ii. **MethQTL calling**, i.e. computing associations between genotype and a DNA methylation state. A two-stage approach is employed: (i) Define CpG correlation blocks as groups of CpGs that are highly correlated across the samples to mimic DNA methylation haplotypes. (ii) From each of these correlation blocks, a tag-CpG is selected as a representative of the block and associations are computed with all SNPs up to a given distance using either a linear modeling strategy or using external software tools (e.g., *fastQTL* [27]). All SNP-CpG pairs that have a P-value below a given cutoff are returned.

#### Data import and preprocessing

##### DNA methylation data

For DNA methylation data, we use the *RnBeads* software package for data handling and processing. *RnBeads* supports most DNA methylation assays yielding single-CpG methylation calls, including whole-genome/reduced-representation bisulfite sequencing (RRBS/WGBS) and the Illumina microarray series. Microarray data can be provided as raw intensity data (IDAT files) and is checked for data quality using *RnBeads’* reporting functionality. Further processing steps, such as CpG and sample filtering (e.g., SNP removal, cross-reactive site removal) and data normalization, can be performed within *RnBeads*. Although we recommend *RnBeads* for data handling, *MAGAR* supports the output of further data processing tools if they provide single-CpG methylation calls.

##### Genotyping data

*MAGAR* accepts microarray and sequencing data as input. Sequencing data has to be preprocessed using genotyping pipelines [55] and converted into a format that is readable through *PLINK* (e.g., VCF files). For microarray data, *MAGAR* supports raw intensity data files as input and computes genotype calls through the *CRLMM* R-package [34,35]. As an optional step, genotyping data can be imputed using the Michigan Imputation Server [54].

Additional data processing, such as filtering SNPs with many missing values or Hardy-Weinberg equilibrium filtering, are conducted through *PLINK*.

#### MethQTL calling

MethQTL calling within *MAGAR* follows a two-stage workflow (**Figure 2A**):

i. CpGs with highly correlated methylation states across the samples are grouped to form CpG-correlation blocks.
ii. A tag-CpG per CpG-correlation block is associated with all SNPs in a given genomic distance to compute associations between SNP genotypes and DNA methylation states.

We elaborate on the two stages in more details below.

##### Correlation block calling

To compute distinct CpG correlation blocks, i.e., groups of CpGs that exhibit high correlations of their methylation states across the samples, from a DNA methylation data matrix we developed a four-step framework:

1. To obtain a similarity matrix, compute the (Pearson) correlation coefficients between the DNA methylation states of any pair of CpGs across the samples using the *bigstatsR* R-package [56] for each chromosome separately. Similarities of two CpGs with correlation lower than 0.2 (package parameter: `cluster.cor.threshold`) are set to zero. Since matrices can grow too large to fit into main memory of standard machines, the CpGs are split per chromosome into equally sized smaller groups until a maximum number of CpGs per computation is achieved (here 40,000 CpGs, parameter: `max.cpgs`).
2. Weight the similarities according to the genomic distance between any CpG and the remaining CpGs using a Gaussian centered at the CpG of interest with standard deviation 3,000 bp (parameter: `standard.deviation.gauss`). Additionally, the similarity between any pair of CpGs further apart than 500 kb is set to zero (parameter: `absolute.distance.cutoff`). Optionally, functional annotations such as the Ensembl Regulatory Build [42] or DNA methylation-based segmentation [57] can be used to re-define the similarities.
3. Construct the associated weighted graph from the similarity matrix, where the edge weights correspond to the similarities between the two CpGs.
4. Employ Louvain clustering [58] using the *igraph* R-package [59] on the weighted graph to obtain clusters of CpGs that are highly correlated. The obtained clusters are defined as the CpG correlation blocks.

The parameters presented here are available as package options to the user. The default parameters have been evaluated using simulations for EPIC and 450k data (**Supplementary Text, Supplementary Figure 2**).

##### Associating SNPs with CpG-correlation blocks

To determine whether the DNA methylation state of a CpG-correlation block is associated with the SNP, we first determine a tag-CpG per correlation block as the medoid of all CpGs in the correlation block. To compute the medoid CpG, we compute the median for each CpG in the correlation block across the samples. Then, we select the CpG that is the median of the vector of medians across the samples as the tag-CpG. Alternative tag-CpG selection methods are available through the package parameter `representative.cpg.computation`. In the next step, all SNPs closer than 500 kb to the tag-CpG are considered and a univariate, least squares regression (lm R function) model is created using the genotypes (encoded as 0=homozygous reference/major allele, 1=heterozygous, 2=homozygous alternative/minor allele) as the features and the CpG methylation state as the response. Further covariates can be included as additional inputs into the linear model. Alternatively, *fastQTL* [27] can be used to compute associations between tag-CpGs and SNPs. The obtained P-values and slopes (referred to as effect sizes or beta in this work) of the linear model are used for further analysis.

#### Package options and modularity

*MAGAR* is a modular software package that allows for easy integration with additional tools. Different variants of the analysis can be specified by the package’s rich option settings. For instance, CpG correlation blocks depend on various parameters including the correlation threshold between two CpGs or the standard deviation of the Gaussian distribution. We used simulation experiments to determine reasonable default parameter settings for the most widely used technologies 450k array, EPIC array, and bisulfite sequencing (**Supplementary Text, Supplementary Figure 2**). However, the option setting can be tailored to the data set at hand. CpG correlation block calling can be deactivated, resulting in the analysis scheme implemented by most published methQTL studies, i.e., associating each CpG with a SNP individually. Additionally, *MAGAR* allows for setting the parameters of the different software tools that are internally used (e.g., *RnBeads*, *PLINK*). To facilitate analyses of large-scale data sets, *MAGAR* supports multi-core processing and automatic distribution of jobs across the nodes of a high performance computing (HPC) cluster. *MAGAR* comes with different export options, including a direct export into the format accepted by GWAS-MAP (see section “Determining tissue-specific methQTLs”). *MAGAR* is publicly available from GitHub (https://github.com/MPIIComputationalEpigenetics/MAGAR).

### Data sets

The data sets used throughout this study have been generated in the context of the SYSCID project (http://syscid.eu/). The CEDAR (Correlated Expression Disease Association Research) [60] cohort data set comprises 164 individuals and we had microarray-based genotyping data available for 163 individuals as described earlier [60]. More specifically, healthy individuals were recruited at the University Hospital in Liège and bowel biopsies as well as blood draws were collected. The biopsies were obtained from rectum (RE) and ileum (IL), and blood cells were FACS sorted into CD4-positive T-cells and CD19-positive B-cells. We used this data set as the discovery cohort within this study. In addition, we used a second data set from the CEDAR cohort comprising additional 197 donors (16 overlapping with the earlier ones) including transverse colon biopsies (n=191) and CD14-positive monocytes (n=192) as a validation cohort.

### DNA methylation profiling

DNA methylation profiling of the samples in the CEDAR cohort was performed using the Illumina EPIC array. Per sample 500 ng of genomic DNA were bisulfite converted using the EZ-96 DNA methylation Gold Kit (Zymo research, Irvine, USA) according to the kit’s manual, except that the final elution volume was reduced to 12 μl. Then, four μl of bisulfite converted DNA was used to run on an Infinium Methylation EPIC array (Illumina, San Diego, USA) according to the manufacturer’s protocol.

DNA methylation data for the validation cohort was generated earlier using the Illumina 450k microarray according to the standard protocol. Due to the small overlap of donors from the EPIC and 450k data set and due to the reduced number of CpGs available on the 450k array, we decided to analyze the data sets separately.

### Genotyping microarrays

Genotyping of the CEDAR cohort has been performed as described earlier [60]. Additional 23 donors have been genotyped using the Illumina Infinium OmniExpress-24v1.3 microarray at the GIGA-Institute Genomics core facility.

### *MAGAR* analysis of the CEDAR cohort

#### DNA methylation data

We used *MAGAR*, which internally uses *RnBeads*, for processing raw IDAT files obtained on the CEDAR cohort samples. A subset (13 B-cell samples, 1 T-cell sample) was removed from the discovery cohort, since the samples exhibited substantially lower technical quality. CpGs were filtered for annotated SNPs in dbSNP [61], for sites on the sex chromosomes, and for potentially cross-reactive sites [62]. Further quality-based filtering of CpGs was conducted using *RnBeads’* Greedycut algorithm [29]. Data was normalized using the “dasen” method from the *wateRmelon* R-package [63]. As outcome of the filtering procedure, 659,464 CpGs were retained for the analysis. The immune cell infiltration was estimated using the LUMP algorithm [32] based on 44 CpGs that are particularly hypomethylated in immune cells, 34 of which were available in the CEDAR data set. For the validation cohort (450k), we used analogous processing options, removed one sample from the 383 samples due to lower technical quality, and retained 353,388 from the 485,777 CpGs available on the microarray.

#### Genotyping data

Genotyping microarray data was imported into *MAGAR* and genotypes called using the KRLMM algorithm implemented in the *CRLMM* R-package [34,35] using default parameters. Genotypes were imputed using the Michigan Imputation Server [54] using *Minimac4* and the following parameters: Reference panel: “hrc-r1.1”, phasing method: “shapeit”, population: “eur”. Imputation was performed for all 163 unique donors simultaneously and the outcome of the procedure yielded 39,127,678 SNPs. Imputed data was exported to *PLINK* [31] for further processing. We filtered for SNPs with a Hardy-Weinberg equilibrium exact test P-value below 0.001, a maximum number of missing values across the samples of 10%, and with minor allele frequency below 5%. Additionally, we removed samples with more than 5% missing genotypes. Finally, 5,436,098 SNPs and all samples were retained.

#### MethQTL analysis

We employed *MAGAR* on an HPC cluster to compute methQTLs for each of the tissues/cell types of the discovery data set independently (**Figure 3B**). Notably, we used sex (categorical), age (continuous), BMI (continuous), smoking state (categorical), alcohol intake (categorical), ethnicity (categorical), and the first two principal components (continuous) computed on the genotype data as covariates in the analysis. *MAGAR* returns a table of methQTL summary statistics (i.e., slope of the regression, standard deviation of the estimate, P-value), which can be further filtered according to a P-value cutoff. Throughout this analysis, we termed methQTLs significant, if they have a P-value below a genome-wide Bonferroni-1.1 adjusted cutoff of 8.65×10^−11^ in the summary statistics returned by *MAGAR*. We computed the P-value cutoff as follows: We identified 82,271, 69,219, 75,779, and 76,109 correlation blocks for T-cells, B-cells, ileum, and rectum samples, respectively. On average, each CpG was tested for association with 1,905 SNPs, which results in:

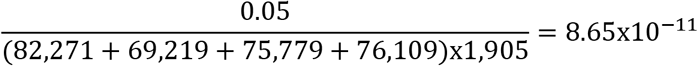

For each CpG that was affected by more than one methQTL, we selected the SNP with the lowest P-value as the lead-SNP.

### Determining tissue-specific methQTLs

To determine whether the effects observed in the four tissues independently were shared across the different samples, we employed colocalization analysis. More specifically, we used Summary-data-based Mendelian Randomization (SMR) and Heterogeneity in Dependent Instruments (HEIDI) analysis [38] implemented in the GWAS-MAP software tool. Briefly, SMR is a statistical test that indicates whether two traits (here CpG methylation states in two tissues) are significantly associated with the same genetic locus. The test is an extension of Mendelian Randomization (MR), which is used to test for a causal relationship between two traits using an instrumental variable. While classical MR requires that the two traits are measured on the same samples, these can be investigated in distinct samples or studies using SMR. The input to the SMR test are methQTL statistics (i.e., P-values, slopes of the regression line) obtained in two scenarios, and it returns a test statistic that indicates whether the effect observed in the two scenarios is significantly associated with the same SNP. Thus, SMR analysis determines whether the same genetic effect leads to the methQTL results that we obtained in the two tissues, but cannot discern pleiotropy (the same SNP influences two traits) from linkage (two highly correlated SNPs influence the traits independently). Thus, for the SNPs that pass the SMR test, we employed the HEIDI test in a second step to test whether the observed effects are likely driven by pleiotropy. Briefly, the HEIDI test utilizes linkage (correlation) information of SNPs from a reference panel to determine whether the observed heterogeneity in the methQTL statistics are more likely caused by linkage than by pleiotropy. By using colocalization analysis through SMR and HEIDI, we were able to determine whether the methQTLs identified in the four tissues/cell types independently were shared or tissue-specific. We employed colocalization analysis for all pairs of tissues/cell types to determine shared methQTLs (**Figure 3B**).

We selected those CpGs for colocalization analysis, which were selected as tag-CpGs in at least two tissues and that had a significant association with a lead-SNP (P-value below 8.65×10^−11^) at least in one tissue. Then, anchoring the analysis on the tissue showing the significant association, we performed the SMR test to detect if the same lead-SNP is associated with the same CpG in any of the other tissues. In case the same lead-SNP was identified in more than one tissue, the tissue/cell type with the lowest P-value was used as the starting point of the SMR analysis. In total, we performed 4,253 SMR tests. The SMR P-values were adjusted for multiple testing using the Benjamini-Hochberg [64] method and we used a P-value cutoff of 0.05. In case the methQTLs measured in two tissues are significant according to the SMR test, this is an indication that the CpG methylation states are significantly correlated with the same SNP in the two tissues. Thus, we use the P-value of the SMR test as an indication of the shared effect of methQTLs in the two tissues.

For CpGs that passed the SMR test, we applied the HEIDI test to discern pleiotropy from linkage. We defined all those pairs of methQTLs with a P-value higher than 0.05 as pleiotropic interactions. The results for a different P-value cutoff (0.001) are shown in **Supplementary Table 3**. The methQTLs that had an SMR test P-value below the cutoff and had a HEIDI test P-value higher than the threshold were defined as shared across the two tissues. The methQTLs shared across all pairwise comparisons according to the colocalization analysis were termed *shared methQTLs*. Additionally, those shared methQTLs with a methQTL P-value below 8.65×10^−11^ in all tissues were termed *common* methQTLs.

The methQTLs that either fail the SMR test or that pass the SMR test, but also pass the HEIDI test were defined as *tissue-specific methQTLs* (**Supplementary Table 3**). Tissue-specificity was defined for each tissue individually. Finally, three classes of methQTLs were defined: tissue-specific, shared, and common methQTLs. SMR and HEIDI analysis was performed using GWAS-MAP (https://www.polyknomics.com/solutions/gwas-map-biomarker-and-intervention-target-discovery-platform).

#### Characterizing tissue-specific and common methQTLs

We merged the methQTLs from the four tissues and compared the effect sizes, P-values, and the distance between the CpG and SNP of the tissue-specific with the methQTLs shared across the tissues. Additionally, we selected different functional annotations of the genome, such as Ensembl genes (version 75), associated promoter regions (defined as 1.5 kb upstream and 0.5 kb downstream of the TSS), and different functional categories according to the Ensembl regulatory build [42]. Then, we overlapped the shared/tissue-specific methQTLs with those annotations using the *GenomicRanges* [65] R-package and computed odds ratios and (one-sided) Fisher-exact test P-values to investigate enrichment towards the functional annotations in comparison to all identified methQTLs. Last, we used the LOLA tool [43] to compute enrichments towards various additional functional annotations from databases such as Cistrome [66], CODEX [67], or ENCODE [68]. In contrast to the annotation enrichment analysis, we performed LOLA enrichment analysis using all CpGs/SNPs that were analyzed as the background for the enrichment.

### Validation of methQTLs

#### Validation using independent data sets

For further validation of the methQTLs identified above, we used 191 transverse colon and 192 monocyte samples from the CEDAR cohort assayed using the Infinium 450k microarray. Genotyping and DNA methylation data was processed analogously to the discovery cohort and methQTLs were called at the P-value cutoff 9.84×10^−6^. We aimed to replicate the 2,508, 696, 1,010, and 868 methQTLs that we identified in the four tissues/cell types and thus computed the P-value cutoff as:

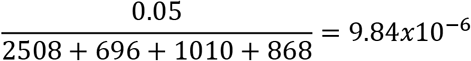

We used sex (categorical), age (continuous), BMI (continuous), smoking (categorical), alcohol intake (categorical), ethnicity (categorical), and the first two principal components (continuous) computed on the genotype data as covariates. The resulting methQTLs were compared with the common and tissue-specific methQTLs detected in the discovery cohort, respectively. Additionally, we obtained methQTL data in tabular form from two studies identifying methQTLs in peripheral blood [12] and fetal brain samples [37], respectively. The two studies identified 52,918 (blood) and 16,811 (fetal brain) methQTLs. We only used unique SNPs with a P-value lower than 8.65×10^−11^ to match our criteria. To determine whether the detected overlap was larger than expected by chance, we used Fisher’s exact test using all SNPs that have been used as input to the methQTL calling as the background set.

#### Validation using local deep sequencing

For validation of the common methQTLs at the *PON1* locus, as well as the shared methQTLs at the *ZNF155*, and *NRG2* loci, we performed local deep sequencing using independent samples from the CEDAR cohort. 500 ng of genomic DNA were bisulfite converted using the EZ-96 DNA methylation Gold Kit (Zymo research, Irvine, USA) according to the kit’s manual. PCRs were set-up in 30 μl reactions using three μl of 10x HotStarTaq buffer (Qiagen, Hilden Germany), 2.4 μl of 10 mM d’NTPs (Fisher Scientific, Pittsburgh, USA), 1.5 μl of 25 mM MgCl_2_ (Qiagen), 0.3 μl each of 10 μM forward and reverse primers (**Table 1**), 0.5 μl of five U/μl HotStarTaq Polymerase (Qiagen), two μl of bisulfite converted DNA, and 20 μl of aqua bidest. PCRs were performed in an ABI Veriti thermo-cycler (Life Technologies, Karlsbad, USA) using the following program: 95°C for 15 min, 40 cycles of 95°C for 30 sec, 1.5 min of 56°C, and one min at 72°C, followed by five min of 72°C and hold at 12°C. PCR products were cleaned up using Agencourt AMPure XP Beads (Beckman Coulter, Brea, USA) and concentration was measured. All amplified products were diluted to four nM and NGS tags were finalized by a second PCR step (five cycles) with primers matching to the NGS compatible tags and carrying a sample-specific barcode (forward 5’-3’: CAAGCAGAAGACGGCATACGAGATXXXXXXGTGACTGGAGTTCAGACGTGTGCTCTTCCGATCT; reverse 5’-3’: AATGATACGGCGACCACCGAGATCTACACXXXXXXTCTTTCCCTACACGACGCTCTTCCGATC; ‘X’s refer to the sample barcode position) followed by a clean-up (AMPure XP). Finally, all samples (set to ten nM) were pooled, loaded on an Illumina MiSeq sequencing machine and sequenced for 2x 300 bp paired-end reads with a MiSeq reagent kit V3 (Illumina) to ca. 10k - 20k fold depth.

**Table1:**
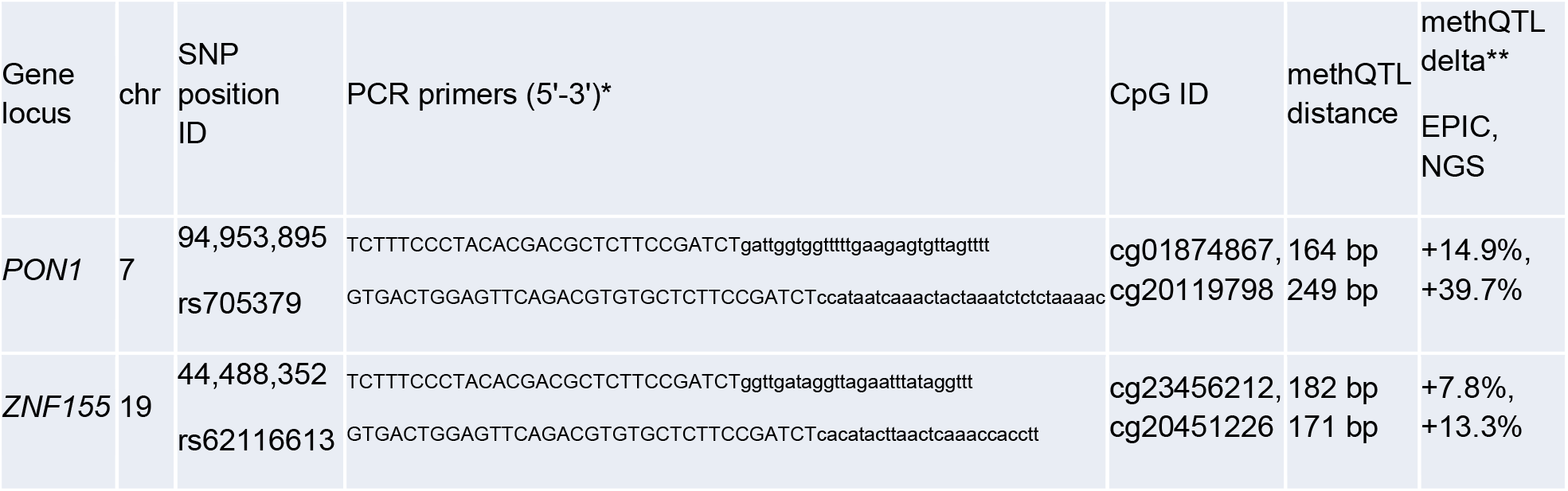
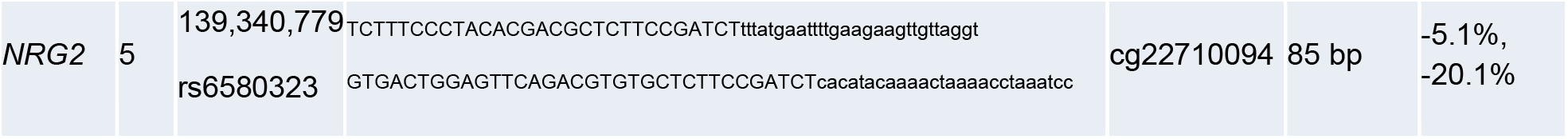
Details on bisulfite amplicons screened in the study. *: Capital letters are NGS compatible tags. ** Absolute methylation change of homozygote minor versus major individuals.

Quality control of the raw data files was performed using the *FastQC* software (https://www.bioinformatics.babraham.ac.uk/projects/fastqc/). Adaptor trimming and filtering for excluding low quality bases was conducted through *cutadapt* [69] and *Trim Galore*! (https://www.bioinformatics.babraham.ac.uk/projects/trim_galore/). Paired reads were joined with the *FLASh* tool [70]. Next, reads were sorted in a two-step procedure by (i) the NGS barcode adaptors to assign samples to identifiers and (ii) the initial 15 bp to assign data to the amplicons. Subsequently, the sorted data was input to *BiQAnalyzer HT* [71] using the following settings: ‘analyzed methylation context’ was set to “C”, ‘minimal sequence identity’ was set to 0.9, and ‘minimal conversion rate’ was set to 0.95. The filtered high-quality reads were used to compute methylation levels of the respective CpGs. Finally, each read was tagged by its base call at the respective SNP position in the amplicon (PON1@173, ZNF155@329, NRG2@255).

## Declarations

### Ethics approval and consent to participate

Not applicable.

### Consent for publication

Not applicable.

### Availability of data and materials

The methylation data used within this publication is accessible at the Data Catalog of the Luxembourg Institute of Health at https://datacatalog.elixir-luxembourg.org/e/study/c7515074-b34a-11eb-9969-acde48001122. Genotyping data has been obtained from the ArrayExpress website (https://www.ebi.ac.uk/arrayexpress) accession number E-MTAB-6666. MAGAR is available through Bioconductor (https://bioconductor.org/packages/release/bioc/html/MAGAR.html).

### Author contributions

M.S. developed MAGAR, generated the figures, and performed the analysis together with G.G., Y.S.A., T.S., and A.N. G.G. generated the DNA methylation data on the samples collected by S.R., E.L., and M.G. G.G. and M.A. performed the local deep amplicon sequencing. D.A. computed cis-regulatory domains on the methylation data. P.A.L., S.R., M.G., Y.S.A., D.A, E.T.D., T.L., and J.W. contributed to the discussion of the results. J.W., T.L., and M.G. jointly supervised the project. M.S. wrote the manuscript with input from all co-authors. All authors approved the final version of the manuscript.

## Acknowledgments

We appreciate the help of Ivan Kuznetsov with data management and analysis and would like to thank Myriam Mni and the GIGA-Institute Genomics core facility for technical assistance. We appreciate the help of the data management team at the Luxembourg Institute of Health, especially by Wei Gu.

## Supplementary tables

**Supplementary Table 1:** MethQTLs identified in the four tissues/cell types using *MAGAR*.

**Supplementary Table 2:** Enrichment P-values according to Fisher’s exact test for validation of the identified methQTLs in independent samples (monocytes, transverse colon) and independent studies (blood and fetal brain).

**Supplementary Table 3:** Results of the colocalization analysis for different P-value cutoffs of the HEIDI test (0.05 and 0.001).

**Supplementary Table 4:** Common methQTLs across the four tissues/cell types.

**Supplementary Table 5:** Tissue-specific methQTLs for the four tissues/cell types.

**Supplementary Table 6:** MethQTLs shared across the tissues/cell types according to the colocalization analysis. The table comprises 1,912 rows and we focus on the 1,470 unique SNPs in the analysis.

**Supplementary Table 7:** GO enrichment analysis results for the shared and tissue-specific methQTLs.

