## Supplementary Text and Figures for "Identification of tissue-specific and common methylation quantitative trait loci in healthy individuals using MAGAR"

##### Supplementary Methods

###### Data Simulation

*MAGAR* is a modular software package that allows multiple parameters to be set for the dataset at hand. We simulated data to determine reasonable default parameters for the two stages of the package independently.

**Validation of correlation block calling** As the first part of the *MAGAR* package, CpGs are grouped together according to their correlation of DNA methylation values across the samples. The process of defining correlation blocks depends on three parameters: the correlation threshold, the standard deviation of the Gaussian distribution, and the absolute distance cutoff. To determine reasonable default values for the parameters of the correlation block calling of the package and to validate that the CpG clustering step is reasonable, we simulated DNA methylation data. More specifically, we simulated methylation data and artificially introduced clusters of highly correlated CpGs into the data. Then, we assessed whether the CpG correlation blocks returned by *MAGAR* reflect the simulated clusters of correlated CpGs. By using different values for the parameters, we were able to assess the parameter setting that best reflects the correlation of CpGs in the simulated data

To simulate methylation data, 1,000 neighboring CpGs were randomly selected from a uniform distribution out of all the CpGs present on the Illumina EPIC array. 1,000 CpGs were selected as a compromise between selecting all CpGs and computational feasibility of executing multiple simulations. We explored different settings for the parameters available in *MAGAR*. First, we explored the influence of the correlation threshold parameter (values tested: 0 to 1, 0.05 steps), which specifies the level of correlation between two CpGs that results in a similarity of zero in the similarity matrix. Second, the standard deviation of the Gaussian distribution (values explored: 2,000 bp to 4,000 bp, 100 bp steps) specifies the width of the Gaussian distribution that penalizes similarities of distant CpGs. The selection of the values for the distance was based on the distances of CpGs on the microarray and should reflect high-to-low penalization of the genomic distance. Last, the absolute distance cutoff (100 kb to 1 Mb, 100 kb steps) sets similarities for long distances between the CpGs to zero. These values were selected, since they also reflect the distance between the SNP and the CpG that defined in the identification of methQTLs (500 kb).

The parameters were tested using 100 simulated datasets per parameter setting. For each of the simulated datasets, we explored the three parameters sequentially and fixed the remaining parameters to the values estimated in the other simulations (starting with 3,000 bp standard deviation and absolute distance cutoff 500 kb for selecting the correlation threshold). Methylation data was simulated by repeatedly drawing from a beta-binomial distribution with success probability 0.4 and over-dispersion parameter 0.1 in order to reflect the typical bimodal distribution of DNA methylation data. For each simulated dataset, we selected the number of clusters randomly between 200 and 500, while choosing the cluster size individually for each cluster between one and ten CpGs. These values should reflect clusters of many CpGs (such as those expected at CpG islands), but also clusters with only few CpGs and were motivated from the distribution of CpGs on the EPIC array. The clusters had identical methylation patterns across the CpGs in the cluster and across the samples. We introduced a Gaussian error for each CpG individually (standard deviation 0.05) to introduce noise into the clusters, which is motivated from our experience on DNA methylation data and the technical noise found on the EPIC array.

To asses *MAGAR's* performance, we executed its first stage and compared the number of expected clusters (i.e., the randomly selected number of clusters) with the number of clusters returned by *MAGAR*. For estimating the parameters for 450k data, we exclusively used CpGs present on the 450k array and for bisulfite sequencing data we used all CpGs available in the human genome reference version “hg19”.

**Validation of methQTL calling** To validate the methQTL calling stage of *MAGAR*, we first generated methylation data as described above. Next, we randomly selected 2,000 SNPs that are located more closely than 500 kb from the CpGs selected. We selected more SNPs than CpGs, since microarray-based systems for genotyping cover more SNPs than DNA methylation microarrays cover CpGs. For those SNPs, we drew the minor allele frequency from a negative binomial distribution (parameter success probability: 0.4) and set the alleles accordingly. We selected the negative binomial distribution, since it most closely reflected the distribution of genotypes in our experimental data. SNP genotypes (α) were encoded as 0=homozygote reference allele, 1=heterozygote, and 2=homozygote alternative allele similar to the standard encoding of *MAGAR*. For each of the 100 simulated experiments that we conducted, we introduced 100 interactions between the genotype of a SNP and the DNA methylation state of a CpG into the data using a randomly selected effect size τ (drawn from a normal distribution with mean 0.2 and standard deviation 0.05). We decided to include only a small number of methQTLs in order to have only few interactions between SNPs and CpGs as we would expect in real data. The sign of the effect size τ was randomly selected as positive or negative, respectively. Similar to the simulation above, we introduced a Gaussian error ϵ into the DNA methylation data. The DNA methylation state β was modified according to the linear model:

$$\beta_{CpG}^{new} = \beta_{CpG}^{old} + \alpha_{SNP} * \tau + \varepsilon$$

We then computed sensitivity and specificity for the CpGs and SNPs independently to assess whether the package successfully identified methQTLs.

#### Supplementary Results

##### Simulated data reveals reasonable default parameters for *MAGAR*

*MAGAR* is a flexible software package that allows users to adapt the analysis to their dataset through different option settings. We determined *MAGAR's* default parameter setting using a simulation strategy. More specifically, we estimated the default parameters for the correlation block calling (**Supplementary Figure 2A-C**) and determined whether methQTLs are reliably detected in the methQTL calling stage (**Supplementary Figure 2D**). We selected the parameters such that the number of clusters (i.e., CpG correlation blocks) returned by *MAGAR* matches the number of clusters that have been simulated. The simulation experiments returned 0.2 as a reasonable parameter for the correlation threshold, which determines at which level of correlation between the two CpGs an edge is removed from the similarity graph. Additionally, 3,000 bp was selected as the standard deviation of the Gaussian distribution, which weights the similarity between two CpGs according to the genomic distance. Notably, higher values for the parameter would more closely reflect the number of simulated clusters in the data. We decided to fix the parameter at 3,000 bp, since we expect that generating fewer cluster (i.e., more singletons) will not have a negative influence on the identified methQTLs. Lastly, we found that the distance cutoff only mildly influences the number of clusters generated, and we determined 500 kb as the distance at which a connection between two CpGs in the graph is removed. This value also matches the maximum distance between the SNP and the CpG that we selected.

To validate whether methQTLs are reliably detected using our package, we artificially introduced interactions between SNPs and CpGs into our simulations. We found high sensitivity and specificity for methQTLs to be detected by *MAGAR* across the 100 simulated datasets that we generated (**Supplementary Figure 2D**). Notably, *MAGAR* was designed to detect reliable methQTLs with few false positive results. Thus, the focus of *MAGAR* is on specificity rather than on sensitivity, but the user can tradeoff between sensitivity and specificity through the p-value cutoff.

#### Correlation blocks are cell-type-specific and largely overlapping with cis-regulatory domains

Although we found that most correlation blocks and their respective tag-CpGs were tissue-dependent, some of the correlation blocks, which we computed for each of the tissues independently, were shared across multiple cell types (**Supplementary Figure 7**). Notably, the overlap between tag-CpGs across the different tissues/cell types was larger than between the correlation blocks indicating that only parts of the correlation blocks were distinct for the different data sets. Additionally, we found that the CpG correlation blocks overlap to a large fraction with cis-regulatory domains (CRDs)[1] identified from DNA methylation data for each tissue/cell type independently (**Supplementary Figure 8**).

### Supplementary Figures

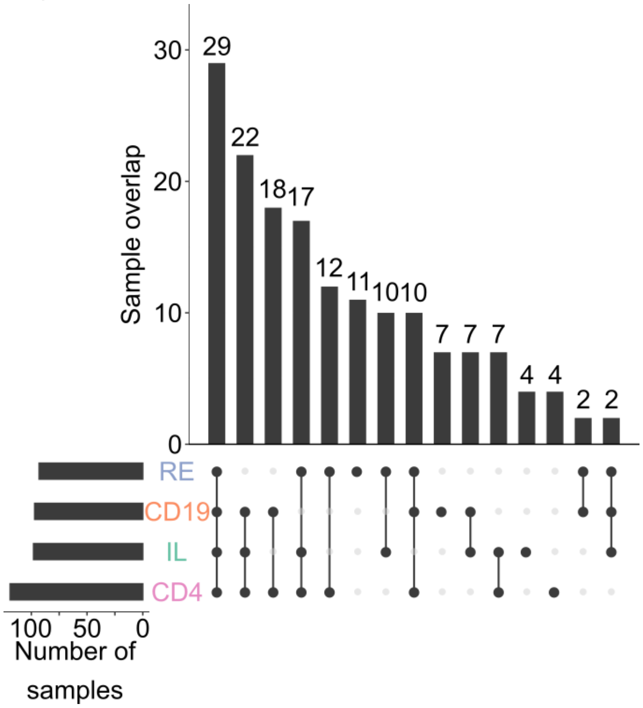


**Supplementary Figure 1: Donor overlap for the samples in the discovery cohort.** UpSetPlot[2] for samples available in the different tissues/cell types that have been assayed in the CEDAR cohort discovery dataset.

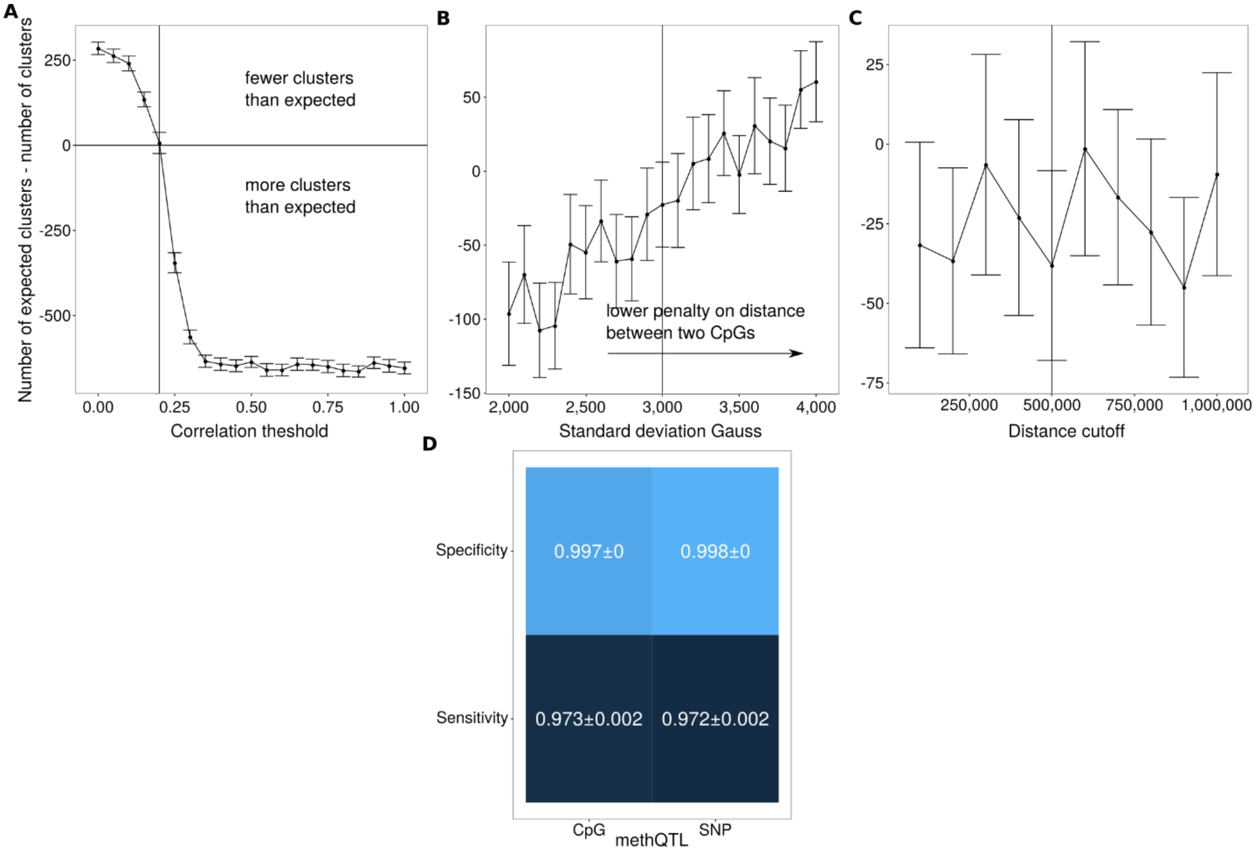


**Supplementary Figure 2: Validating *MAGAR* and its parameters using simulated data**. **A:** Difference between the number of expected clusters and the number of clusters returned by the package in comparison to the correlation threshold parameter. Higher values on the y-axis indicate that the clusters/correlation blocks generated by the package are too large. The solid line indicates the selected value of the parameter for EPIC data. The error bars indicate two times the standard error computed across the 100 simulated data sets per parameter setting. Influence of the standard deviation of the Gauss distribution (**B**) and the absolute distance cutoff (**C**) on the number of clusters. **D**: Sensitivity and specificity of *MAGAR's* methQTL calling in simulated data. Shown are the mean and the standard error across the 100 simulated data sets.

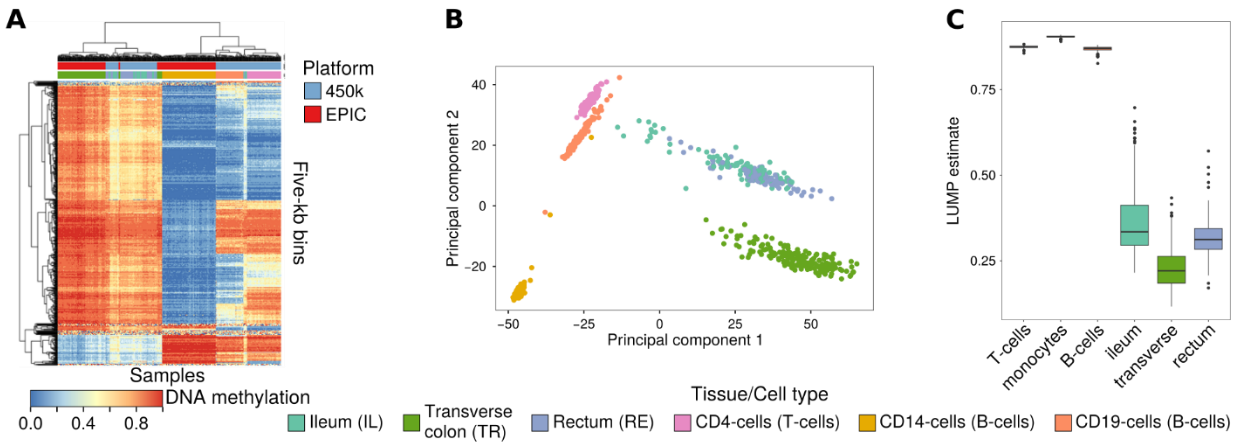


**Supplementary Figure 3: Joint description of the validation and discovery data set. A:** Heatmap for methylation states of genome-wide bins (5 kb) for a combination of the discovery data set (EPIC) and the validation data set (450k) from the CEDAR cohort. Analysis was restricted to the intersection between the 450k and EPIC CpGs. **B:** PCA plot for CpG-wise DNA methylation beta-values for the different samples in the combined EPIC/450k dataset. **C:** LUMP estimates of the overall immune cell content stratified according to the six different tissues/cell types.

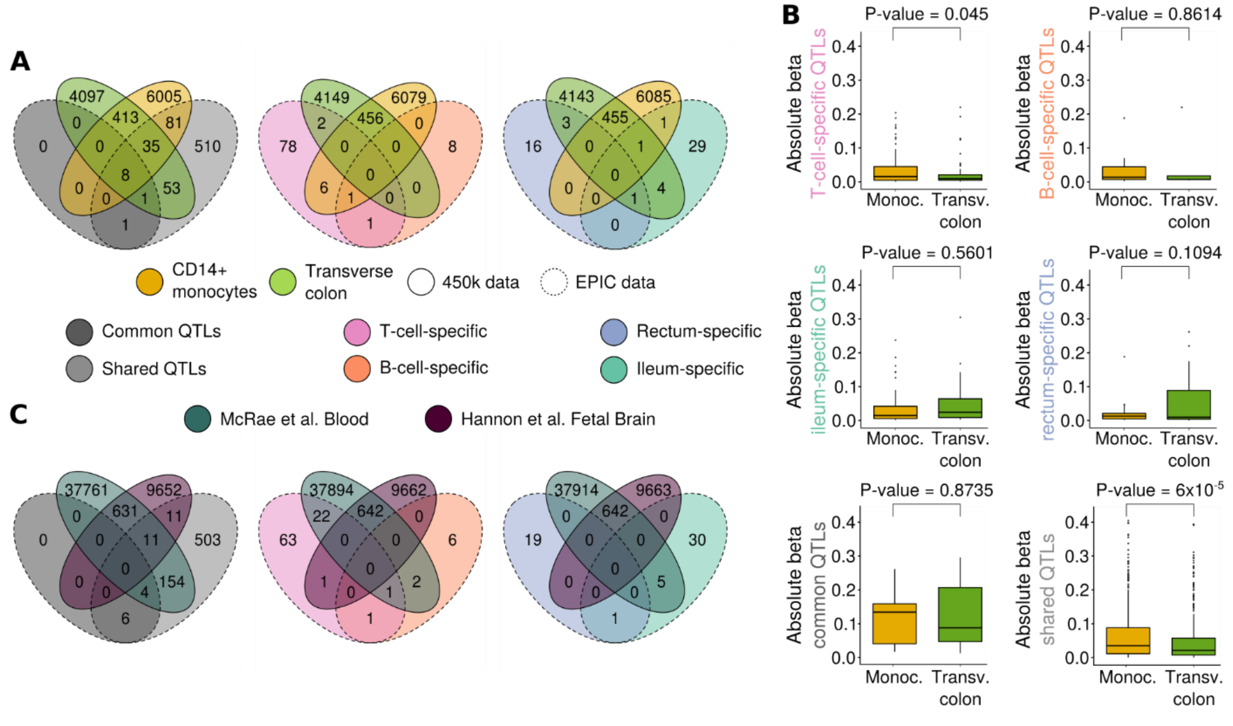


**Supplementary Figure 4: Validation of tissue-specific and common methQTLs in the validation cohort and in two independent studies. A:** Overlap of tissue-specific, common, and shared methQTLs with methQTLs identified in CD14-positive monocytes and transverse colon samples assayed with the Infinium 450k microarray. **B:** Effect size comparison of the T-cell-specific, B-cell-specific, shared, and common methQTLs in CD14-positive monocytes and transverse colon samples. **C:** Overlap of tissue-specific, shared, and common methQTLs with published methQTL studies in blood and fetal brain samples.

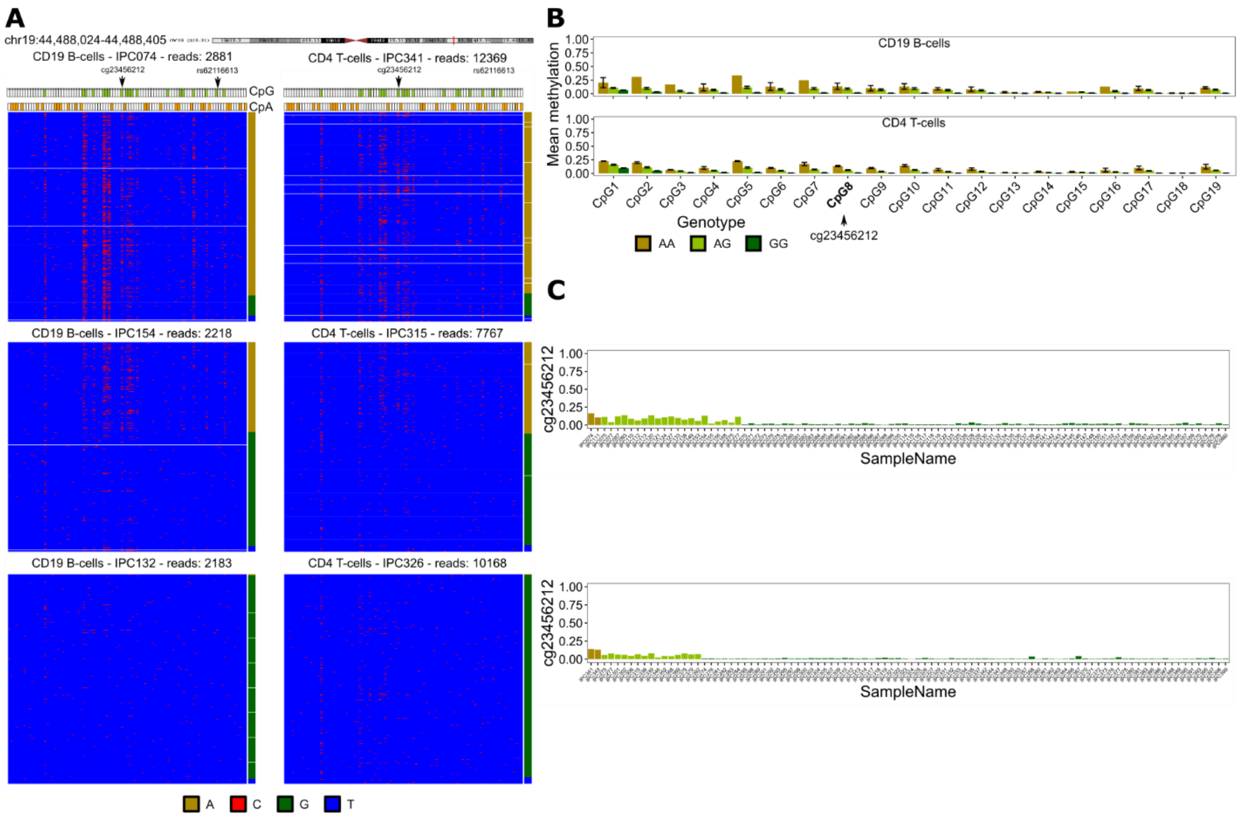


**Supplementary Figure 5: Validation of methQTL at *ZNF155* locus using ultra-deep bisulfite sequencing. A:** Bisulfite sequencing read pattern maps for three individuals with genotypes homozygous reference allele (AA), heterozygous (AG), and homozygous alternative allele (GG) for B-cells and T-cells, respectively. Each line is a sequencing read, where the red color indicates a cytosine, i.e. a methylated cytosine before bisulfite conversion, and blue a thymine, i.e., an unmethylated cytosine before bisulfite conversion. All cytosines within the amplicon are shown in the pattern map, and the CpG and CpA dinucleotides are marked. The genotype of rs62116613 per sequencing read is indicated on the right. Shown is the *ZNF155* locus (shared methQTL) at chr19:44,488,024-44,488,405 (hg19). **B:** Average DNA methylation levels across all the samples of the same genotype and standard deviations across the samples. The barplots are shown for all 19 CpGs present in the amplicon. **C:** Average DNA methylation levels across all sequencing reads per sample for the CpG that was associated with the SNP genotype in the microarray data analysis for B-cells and T-cells.

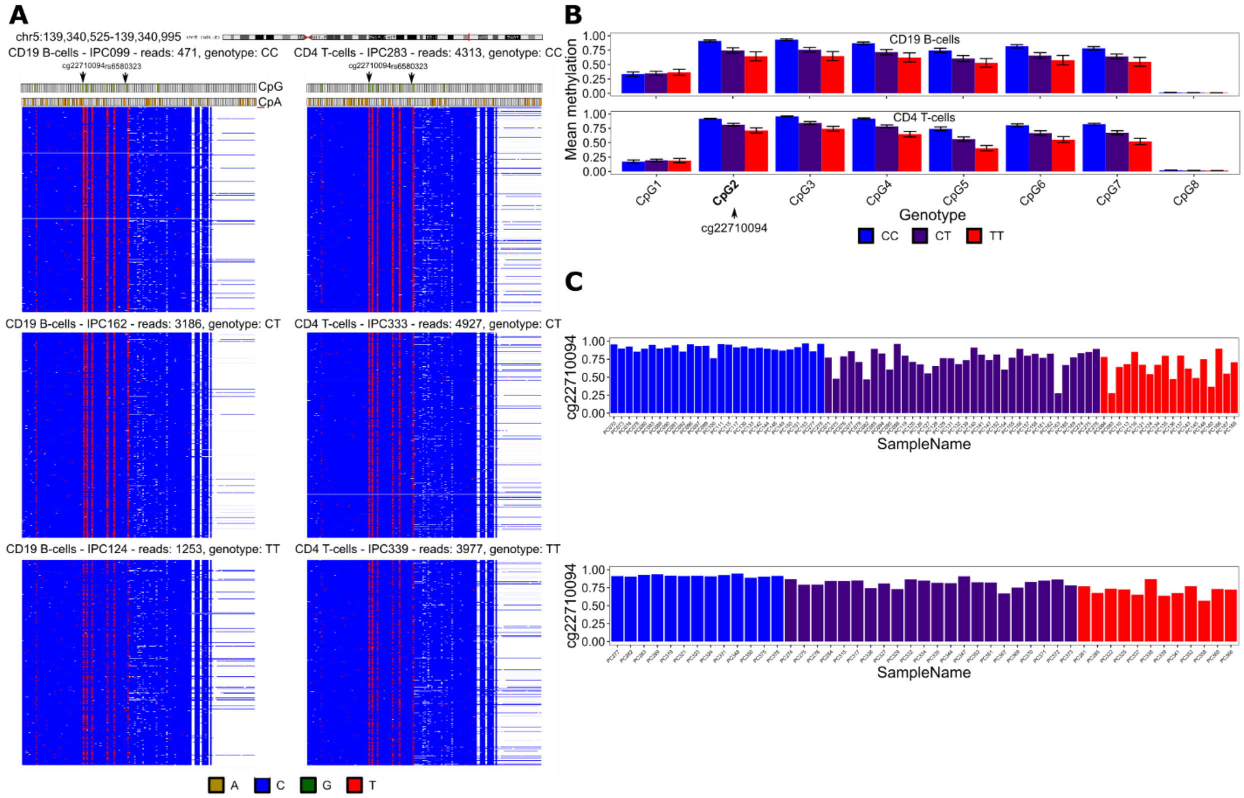


**Supplementary Figure 6: Validation of methQTL at *NRG2* locus using ultra-deep bisulfite sequencing. A:** Bisulfite sequencing read pattern maps for three individuals with genotypes homozygous reference allele (CC), heterozygous (CT), and homozygous alternative allele (TT) for B-cells and T-cells, respectively. Each line is a sequencing read, where the red color indicates a cytosine, i.e., a methylated cytosine before bisulfite conversion, and blue a thymine, i.e. an unmethylated cytosine before bisulfite conversion. All cytosines within the amplicon are shown in the pattern map, and the CpG and CpA dinucleotides are marked. The genotype of rs6580323 per sequencing read could not be computed, since unmethylated cytosines and thymines cannot be differentiated after bisulfite conversion. Thus, the genotype from the genotyping microarray was used here. Shown is the *NRG2* locus (shared methQTL) at chr5:139,340,525-139,340,995 (hg19). **B:** Average DNA methylation levels across all the samples of the same genotype and standard deviations across the samples. The barplots are shown for all eight CpGs present in the amplicon. **C:** Average DNA methylation levels across all sequencing reads per sample for the CpG that was associated with the SNP genotype in the microarray data analysis for B-cells and T-cells.

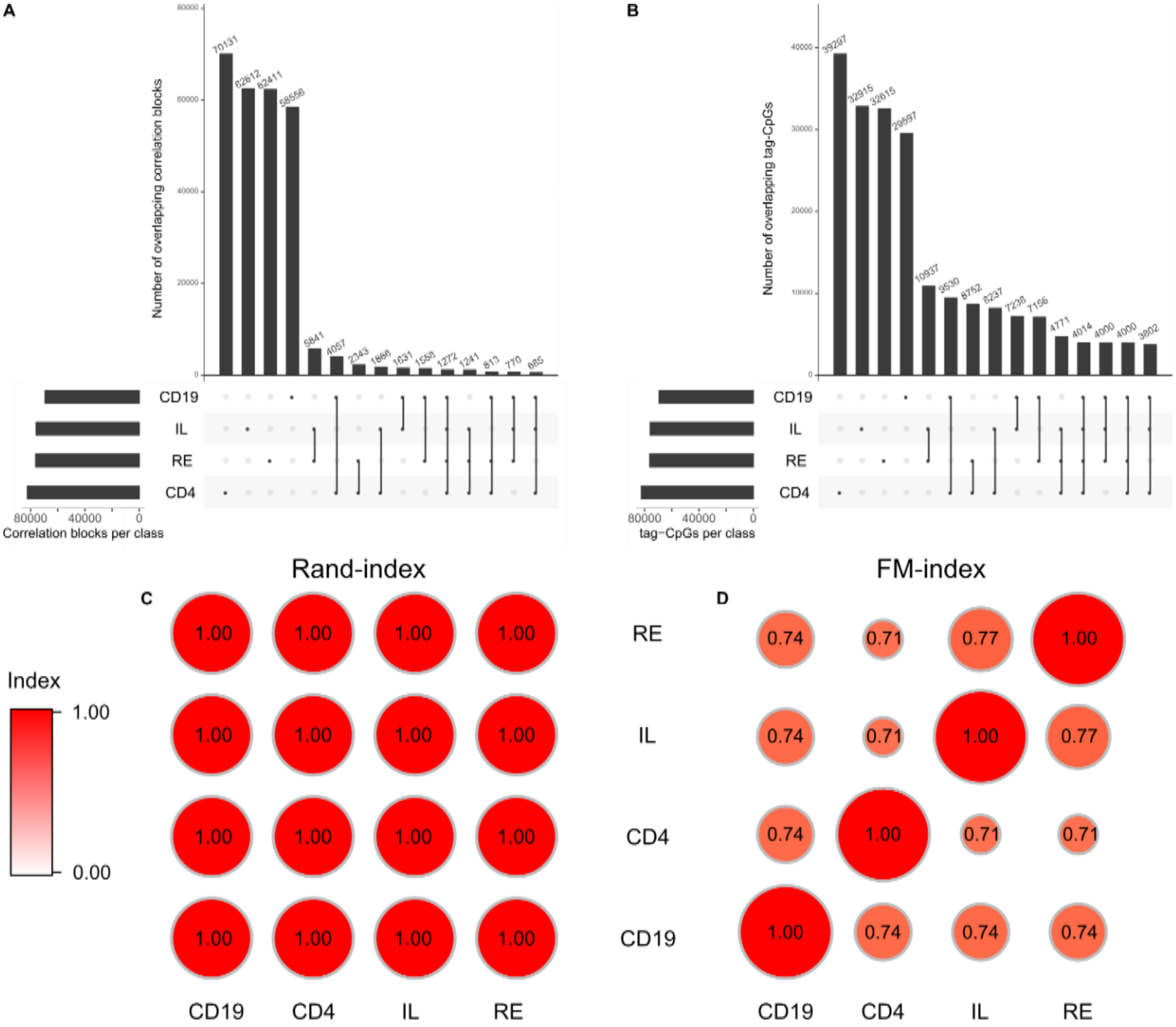


**Supplementary Figure 7: Overlapping correlation blocks (A) and tag-CpGs (B) per cell type/tissue using the parameter setting described in Supplementary Figure 2**. **A**: Overlapping correlation blocks for the four tissues/cell types generated on the EPIC array. Correlation blocks were considered identical if all CpGs in the two correlation blocks defined in the two tissues independently are shared. **B**: Overlapping tag-CpGs for each of the correlation blocks per tissue/cell type. Tag-CpGs were computed for each of the correlation blocks and each of the tissues/cell types independently. **C:** Rand indeces[3] for the different correlation blocks computed across the cell types/tissues. **D:** Fowlkes-Mallows (FM)[4] index for the different correlation blocks computed across the cell types/tissues.

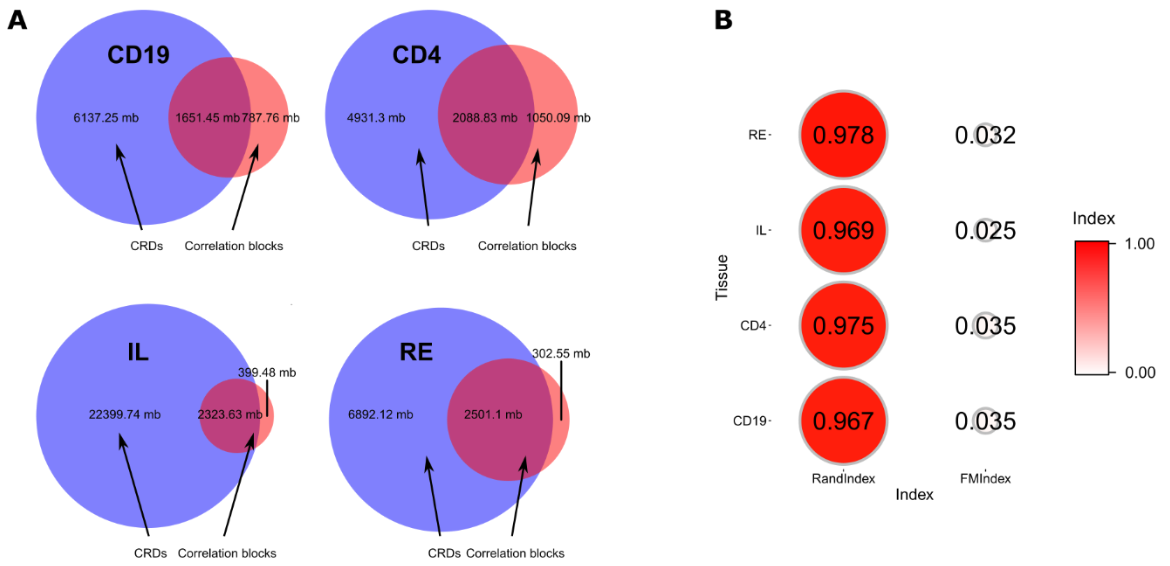


**Supplementary Figure 8: Comparison of correlation blocks with cis-regulatory domains[1]. A:** Genomic overlap of cis-regulatory domains (CRDs) and CpG correlation blocks. The genomic size of the correlation blocks was defined as the distance between the first and the last CpG. **B:** Rand- and Fowlkes-Mallows (FM)-indices comparing the clustering defined by the CpG correlation blocks and the CpGs within the same CRD.
